# Segmental regulation of intestinal motility by colitis and the adaptive immune system in the ileum and colon

**DOI:** 10.1101/2024.04.29.591625

**Authors:** Raquel Gómez Bris, Pilar Rodríguez-Rodríguez, Marina Ortega Zapero, Santiago Ruvira, Angela Saez, Silvia Magdalena Arribas, Jose Maria Gonzalez Granado

## Abstract

Gastrointestinal motility disturbances are common in inflammatory bowel disease (IBD); however, their exact causes remain elusive.

This study explores the motility of various intestinal segments in both healthy and IBD states, focusing on the role of the adaptive immune system. Using a dextran sulfate sodium (DSS)-induced colitis model in mice lacking B and T lymphocytes, we evaluated motility in the ileum and colon using an organ bath system.

In healthy mice, absence of adaptive lymphocytes in the ileum reduces muscarinic receptor sensitivity or increase cholinesterase activity. Colitis increases motility, intensifying the intensity and frequency of spontaneous contractions while decreasing responsiveness to cholinergic stimuli.

In the proximal colon, healthy mice lacking adaptive immune system exhibit increased contractile capacity and frequency, along with reduced muscarinic receptor sensitivity or increased cholinesterase activity. Conversely, colitis diminishes contractile capacity regardless of genotype, while recovery increases frequency of spontaneous contractions.

In the mid-colon during colitis, healthy mice lacking adaptive immune system exhibit reduced muscarinic receptor sensitivity or increased cholinesterase activity, while the absence of adaptive lymphocytes during colitis exacerbates both spontaneous and stimuli-induced contractions. Finally, in the distal colon, the adaptive immune system enhances stimuli-induced contractility in health and reduces contractility and enhances muscarinic responses during colitis.

Overall, intestinal motility in both the ileum and colon is finely regulated, with the adaptive immune system playing a crucial role. These findings contribute to our understanding of IBD pathology, emphasizing the importance of investigating gastrointestinal motility in IBD research.

## Introduction

The gastrointestinal tract is divided into the small and large intestines, hosting a dense population of commensal microbiota ^1^. This microbial community displays spatial heterogeneity along the proximal-distal axis, contributing to the establishment of a highly specialized organ with unique functions depending on its position along this axis ^2-4^. While the high degree of compartmentalization in the small intestine is well-defined, the extent of molecular regionalization in the colon is starting to be better understood ^5^.

The intestine depends on continuous regeneration of the intestinal epithelium and control of the immune system to maintain homeostasis. Disruption of these processes may result in the onset of chronic intestinal conditions like inflammatory bowel disease (IBD). Consequently, following injury, the intestine must promptly adapt to minimize tissue damage, facilitate tissue regeneration and promote recovery. To achieve this, immune cells are recruited or expanded *in situ* to protect the host from invading pathogens and to coordinate the healing process by emitting resolving signals ^6^.

IBD is a chronic inflammatory disorder of the intestine that manifests as Ulcerative Colitis (UC) and Crohn’s Disease (CD). It arises from an inappropriate host immune response to commensal bacteria in genetically susceptible individuals. The overall pathophysiology of the disease is well-understood and comprises several interconnected processes: environmental influences can alter the composition of the gut microbiota, impacting mucosal barrier function and leading to epithelial alterations. This can ultimately result in inflammation in the lamina propria^7-10^. When the integrity of the intestinal epithelium is compromised, certain microbiota components can cross into the lamina propria, inducing the activation of dendritic cells (DCs) and macrophages. This cascade of innate immune cells activation culminates in the recruitment of CD4+ T cells to the intestinal tissue. Individuals with IBD typically exhibit an increased presence of proinflammatory T helper (Th) cells, specifically Th1 and Th17, coupled with insufficient numbers of immunosuppressive cells, such as regulatory T cells (Tregs). The imbalance towards pro-inflammatory T cells guides the function of cells with an innate immune role, such as epithelial cells, fibroblasts, and phagocytes, leading to a persistent hyperresponsiveness to microbial antigens, and consequent tissue injury, perpetuating chronic intestinal inflammation ^7-9^.

While most research has focused on investigating and treating the inflammation and histological alterations within the digestive tract of individuals with IBD, there is a scarcity of studies examining the impact of IBD on gastrointestinal motility. The digestive tract exhibits diverse motility patterns that are regulated by complementary and overlapping control systems, including the enteric and autonomic nervous systems, smooth muscle cells (SMCs), and interstitial cells of Cajal (ICCs) ^11,12^. These patterns differ in intensity and rhythm across different parts of the gastrointestinal system ^13^.

Interestingly, several factors implicated in the development of IBD are believed to be associated with gastrointestinal motor abnormalities ^14^. Furthermore, it has been reported that certain motility disorders, previously labelled as “idiopathic”, are indeed attributable to low-grade inflammation. Examples of such gastrointestinal disorders include esophageal achalasia, functional dyspepsia, irritable bowel syndrome (IBS) and chronic constipation ^14^. Moreover, alterations in the enteric nervous system (ENS) have been observed in patients with IBD, which affect the typical motor function and visceral reflexes of the gastrointestinal tract ^14^. However, the precise impact of the immune system on intestinal motility remains poorly understood. Investigations conducted in various mouse models have yielded conflicting results, showing both increases and decreases in motility depending on the region of intestine studied or type of insult. Various models of inflammation have shown a decrease in motility ^13^. Conversely, infection has been shown to cause hypercontractility, which is associated with increased Interleukin-9 and dysregulation of Th and Treg ^15^. The primary objective of this study was to evaluate the role of the adaptive immune system in the physiopathology of IBD, specifically in relation to intestinal motility. Additionally, the study aimed to identify any differences between the small and large intestine contractility in both homeostasis and colitis conditions.

## Material and Methods

### Mice

Experiments were conducted on male mice aged 8-14 weeks, including both wild-type (WT) C57BL/6J mice and C57BL/6J Rag1^-/-^ mice lacking B and T lymphocytes ^16^. The mice were bred under specific pathogen-free (SPF) conditions at the Centro Nacional de Investigaciones Cardiovasculares (CNIC) in accordance with the sanitary recommendations of the Federation of European Laboratory Animal Science Associations (FELASA). The experimental procedures involving animals were approved by the Fundación Centro Nacional de Investigaciones Cardiovasculares Carlos III (CNIC), Univesidad Autónoma de Madrid (UAM) and the Comunidad Autónoma de Madrid, in accordance with Spanish and European guidelines.

### DSS Colitis model

To induce colitis as previously described ^17^, mice were administered 2% (w/v) dextran sulfate sodium salt (DSS) (Alfa Aesar) in their drinking water for 5 days. This was followed by a recovery period of 3 to 7 days during which they consumed only water. Mice from each genotype (WT and Rag1^-/-^) were divided into three groups: healthy group (exposed solely to water), colitis group (sacrificed on day 8 post-DSS administration) and recovery group (sacrificed on day 12 post-DSS administration). Throughout the experimental period, daily monitoring of mice body weight was diligently conducted.

### Solutions

KHS (Concentration (mM)): NaCl 115, KCl 4.7, MgSO4*7 H2O 1.2, KH2PO4 1.2, NaHCO3 25, CaCl2 2.5, EDTA 0.01 and glucose 11.1) previously refrigerated and gassed with 5% CO2 and 95% oxygen gas. Calcium-free KHS (0Ca2+) (Concentration (mM)): 115.0 NaCl, 25.0 NaHCO3, 4.7 KCl, 1.2 MgSO4*7 H2O, 1.2 KH2PO4, 11.1 glucose and 10 EGTA). KCl solution (Concentration (mM)): KCl 120, MgSO4*7 H2O 1.2, KH2PO4 1.2, NaHCO3 25, CaCl2 2.5, EDTA 0.01 and glucose 5.55).

### Organ bath

After sacrifice, the entire intestine, from the ileum to the anus, was isolated and preserved in cold Krebs Henseleit solution (KHS). Subsequently. The ileum and colon underwent meticulous cleaning by gently flushing with KHS. The colon was divided into three regions (proximal, middle and distal to the ileum). From each mouse four segments were taken from the ileum, one from the proximal colon, two from the middle colon, and one from the distal colon measuring approximately 5-6 mm. The segments were mounted longitudinally in individual organ bath chambers as follows: a piece of suture thread was tied to each end of the intestine segment; one thread was attached to a fixed metal support and the other end of the intestine was connected to an isometric force transducer ^18^. The chamber contained KHS gassed with a mixture of 95% oxygen and 5% CO_2_ maintaining pH of 7,3 to 7,4. The tissues were stretched to 0,5 g or 1 g for ileum and colon respectively and allowed to equilibrate. Equilibration lasted until the emergence of spontaneous contractions or for a minimum duration of 40 min. After recording the spontaneous contractions, the tissue was exposed to a depolarization solution by replacing the KHS with a 120mM KCl solution. Following the washout period, dose-response curves were studied for muscarinic agonists, one for carbachol (Sigma-Aldrich) and one for acetylcholine (ACh, Sigma-Aldrich), at concentration ranging from 10^-9^ M to 10^-4^ M, with at least 30-min washout period between them. Ach and carbachol stock solutions were prepared in saline ascorbic acid to prevent oxidation. At the end of the experiment, a basal measurement was obtained by using KHS without calcium. All parameters were measured in all mounted segments. The data were recorded and analyzed using LabChart 8 software (ADInstruments Ltd, ^19^).

### Statistical analysis

Statistical analysis and graphics were performed using GraphPad Prism 9. Two-way ANOVA with Šídák’s multiple comparisons test was used for the analysis of weight evolution. Multiple comparisons between all groups were conducted using one-way ANOVA and Tukey post-hoc tests. Student T-Test was used to compare WT and Rag1^-/-^ healthy groups. Maximal spontaneous contraction and KCl contraction were corrected with the length of the segment. For the analysis of the carbachol and Ach curves, we related the contraction of each agonist to the maximal KCl contraction and corrected it to start at 0. We then calculated the area under the curve (AUC), maximal responses and PD2 (-logEC50) to assess the efficacy and sensitivity of the muscarinic receptors. We considered a p value of <0.05 to be significant and <0.1 to be a tendency.

## Results

### Mice administered DSS develop colitis

The study defined six experimental groups based on weight change profiles. These groups included non-treated healthy WT mice (healthy WT), WT mice treated with DSS for 5 days and killed on day 8 (WT colitis) or day 12 (WT recovery), non-treated healthy Rag1^-/-^ mice (healthy Rag1^-/-^), and Rag1^-/-^ mice treated with DSS for 5 days and euthanized on day 8 (Rag1^-/-^ colitis) or day 12 (Rag1^-/-^ recovery). Non-treated WT and Rag1^-/-^ mice exhibited a non-significant slight increase in weight from the beginning of the experiment until day 12, with no significant differences between the two groups (Figure 1 A). Both DSS-treated WT and Rag1^-/-^ mice experienced gradual weight loss until 7-8 days after the start of treatment, indicating the development of colitis. They then regained weight until the end of the experiment on day 12. By day 8, both the DSS-treated WT and Rag1^-/-^ mice experienced significant body weight loss compared to the non-treated WT and Rag1^-/-^ mice. By day 12, the body weights of both the DSS-treated WT and Rag1^-/-^ mice were similar to those of the non-treated WT and Rag1^-/-^ mice. The DSS-treated Rag1^-/-^ mice exhibited less weight loss on day 8 and reached similar weight levels by day 12 compared to the WT mice treated with DSS.

**Figure 1.**
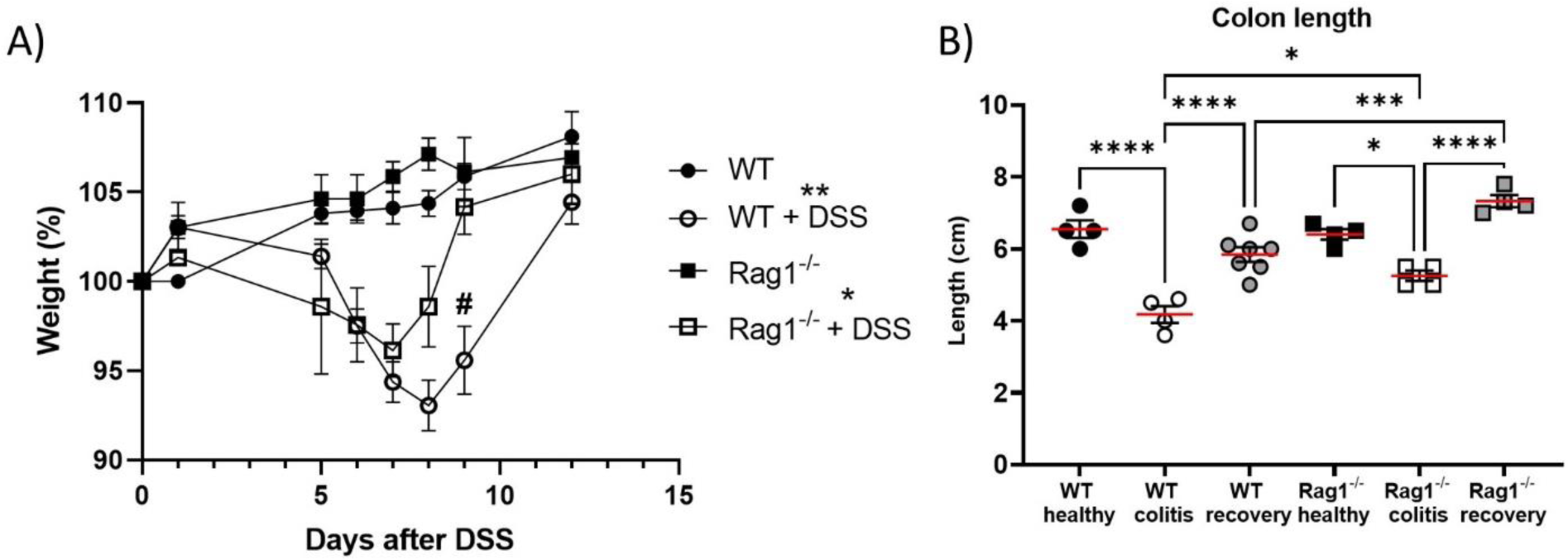
DSS-induced colitis. A) Evolution of mice weight as a percentage relative to day 0. Mice from different groups were weighted throughout the experiment, with day 0 designated as the day when groups receiving DSS were given 2% (w/v) in their drinking water. Mice from non-treated groups were weighted concurrently. Two-way ANOVA analysis (n = 4 - 7), * p < 0.05, ** p < 0.01 within same genotype, # p < 0.05 respect same treatment. B) Colon length measurement. One-way ANOVA analysis (n = 4 - 7), * p < 0.05, *** p < 0.001, **** p < 0.0001.

Upon isolation, the length of the colon was measured (Figure 1 B). The study found that WT and Rag1^-/-^ colitis mice had shorter colons than healthy WT and Rag1^-/-^ mice on day 8, indicating colon inflammation. By day 12, the colon length of recovery WT and Rag1^-/-^ mice was significantly different from that of the WT and Rag1^-/-^ colitis mice, but not from that of the healthy WT and Rag1^-/-^ mice, respectively, indicative of complete recovery. No differences in colon length were observed between the healthy WT and Rag1^-/-^ groups. However, the colon length of the colitis and recovery Rag1^-/-^ mice was greater than that of colitis and recovery WT mice, respectively (Figure 1 B).

These findings suggest that DSS treatment induces colitis in mice, with the notable difference that Rag1^-/-^ mice develop a milder pathology and a recover more effectively, indicating a role for the adaptive immune system in the pathology. The absence of adaptive lymphocytes leads to a less severe phenotype.

### Ileal spontaneous contractions are influenced by the presence of adaptive lymphocytes in acute colitis

In our organ bath experiments, we assessed spontaneous contractions by measuring their maximal contraction and frequency (Figure 2 A). We studied two states of colitis: acute colitis, characterized by a period of weight loss during which we examined mice at their minimum weight by day 8; and recovery, when mice completed the process of weight regain, reaching either the same percentage as healthy groups or at least 100% of the weight from day 0.

**Figure 2.**
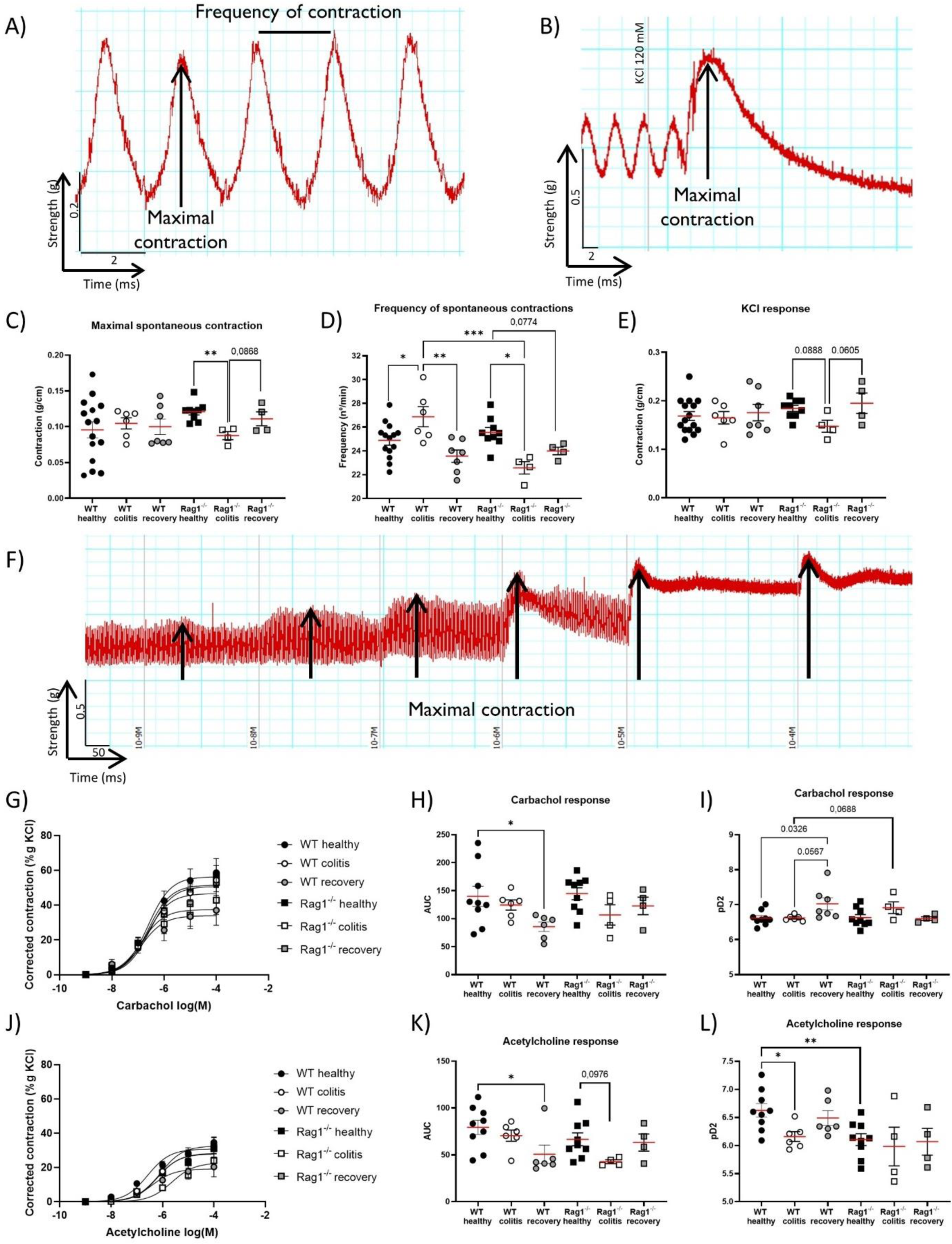
Data analysis of ileum. A) Example of how Maximal contraction and frequency of spontaneous contractions are measured in the recorded data. B) An illustrative example showing the characteristic KCl contraction pattern. The methodology employed for measuring maximal contraction is also outlined. C) Maximal spontaneous contractions. D) Spontaneous contractions frequency. E) Results of KCl maximal contraction. F) Illustrative example depicting the appearance of contraction curves with increasing concentrations and how maximal contraction are measured for each concentration. G-L) Results of carbachol (G-I) and acetylcholine (J-L) contraction curves (G, J), AUC (H, K), and PD2 (I, L). One-way ANOVA (n = 4 - 15) * p < 0.05, ** p < 0.01, *** p < 0.001.

In healthy conditions, no differences in maximal contractions were detected between WT and Rag1^-/-^ mice. In WT ileum, no disparities were observed among the healthy, colitis, and recovery groups. However, in Rag1^-/-^ mice, the colitis group exhibited lower maximal spontaneous contractions compared to healthy and recovery Rag1^-/-^ ileum (Figure 2 C). Regarding the frequency of contractions, there were no differences between genotypes in healthy conditions (Figure 2 D). In WT ileum, acute colitis increased the spontaneous contraction frequency, with this effect returning to similar levels as the healthy conditions in the recovery WT group. Conversely, in Rag1^-/-^ ileums, acute colitis resulted in the opposite trend, with a lower frequency of spontaneous contractions observed in colitis Rag1^-/-^ samples when compared to healthy Rag1^-/-^ mice. Additionally, an incomplete recovery of the frequency of spontaneous contractions was observed in the Rag1^-/-^ recovery group.

Maximum spontaneous contractions and their frequency are induced by acute colitis and reduced in the absence of B and T lymphocytes, giving the primary distinction between genotypes during the acute colitis phase (Figure 2).

### The contractile response of the ileum to stimuli is influenced by colitis and the presence of adaptive lymphocytes

We examined Ileum contractions in response to stimulus by studying the maximal response to a depolarizing potassium solution (Figure 2 B), and the contractile response to the muscarinic agonists carbachol and acetylcholine, which were studied using dose-response curves (Figure 2 F).

In healthy conditions, no significant differences were observed between WT and Rag1^-/-^ mice. In WT mice (Figure 2 E), there were no significant differences in the contractile capacity between healthy individuals and different stages of colitis disease. However, in Rag1^-/-^ mice, ileums from those with active colitis displayed a tendency to decrease in their maximal contractile capacity, which returned to similar levels as healthy Rag1^-/-^ samples in the recovery group. Subsequently, we studied the contractile response to muscarinic agonists, carbachol (Figure 2 G-I) and acetylcholine (Figure 2 J-L). This experimental design allowed us to compare the response to the physiological agonist, acetylcholine (including its action on muscarinic receptors and its degradation by acetylcholinesterase), with that of carbachol, an analogue of acetylcholine which exhibits resistance to degradation by acetylcholinesterase.

In the WT ileum under healthy conditions, the area under the curve (AUC) was larger for carbachol than for acetylcholine, indicating active acetylcholinesterase in ileal preparations under our experimental conditions. This effect was also observed in healthy Rag1^-/-^ samples (Figure 2 H and K). Moreover, we observed that carbachol and Ach responses did not differ between genotypes under healthy conditions. Colitis did not significantly affect responses to both agonists in WT mice, while Rag1^-/-^ presented a reduced response to Ach due to colitis (tendency). A significant reduction in response to both agonists was observed in WT ileums upon recovery when compared to healthy samples, and effect that was abolished in Rag1^-/-^ ileums (Figure 2 H and K).

Regarding the PD2 (negative logarithm of the EC50), in healthy conditions, no significant differences were found in carbachol responses between WT and Rag1^-/-^ mice. However, Rag1^-/-^ mice showed a lower PD2 values to acetylcholine compared to WT, suggesting a lower sensitivity of muscarinic receptors (or more active acetylcholinesterase) in mice lacking acquired immune system. In WT mice with colitis, PD2 values for Ach, but not to carbachol, were also significantly lower compared to healthy conditions (or increased acetylcholinesterase activity induced by DSS), becoming normal upon recovery. This was not observed in Rag1^-/-^ mice, probably due to the fact that they already had a lower basal sensitivity to acetylcholine (or more active basal acetylcholinesterase). However, in mice with colitis, the PD2 value to carbachol tended to be higher in Rag1^-/-^ mice compared to WT, suggesting increased muscarinic receptor sensitivity in mice lacking the adaptive immune system, and in WT recovery group in comparison with healthy and colitis.

In summary, in ileum, the adaptive immunity does not affect the maximal contractile response to stimuli but enhances sensitivity to Ach; colitis also reduces sensitivity to ACh but only in WT mice; and recovery reduces the maximal response to muscarinic agonists.

### Colon intestinal motility exhibits geographic heterogeneity

We further investigated the response in the organ bath system of the mouse colon. To perform a comprehensive study, considering the diverse physiological roles of the different segments of the colon, we divided the entire colon in four parts based on their distance to the cecum: proximal, two middle sections and distal. In the colon segments, we identified a pattern of large spontaneous contractions like those found in the ileum. Additionally, we detected small contractions within these large contractions, and we measured the frequency of both (Figure 3).

**Figure 3.**
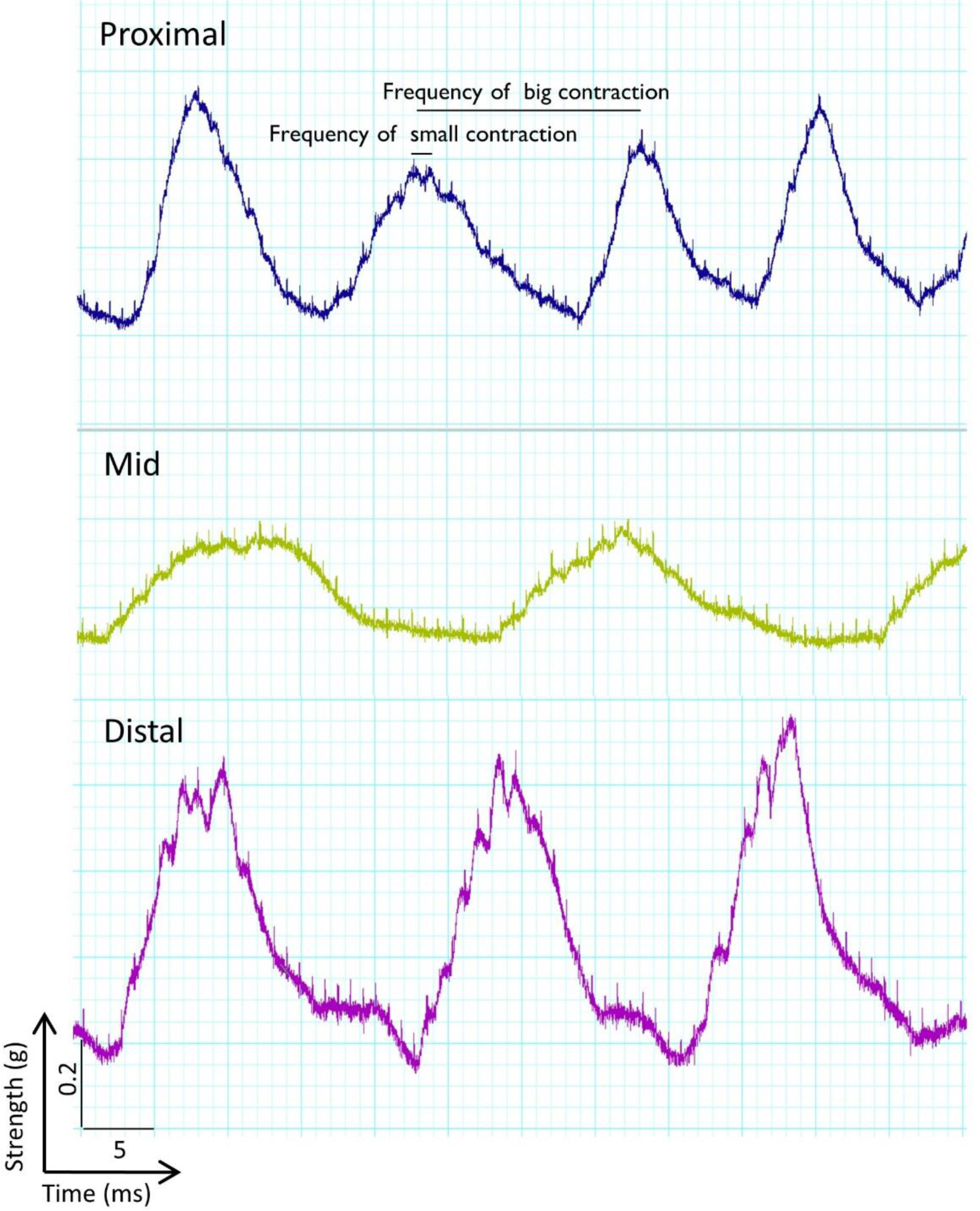
**Example of records obtained in the organ bath system for a proximal, a mid and a distal segment**, illustrating how the frequency of big and small contractions are measured in the recorded data.

When comparing segments from the colon of WT mice (Figure 4), we observed distinct patterns. While distal colon exhibited a greater contractile response of spontaneous contractions compared to the mid and proximal sections (Figure 4 A), proximal colon presented the highest frequency of these spontaneous contractions, particularly small contractions (Figure 4 B, C). Moreover, distal segment was also characterized by a more substantial contractile response to KCl and carbachol, but not ACh compared to mid and proximal segments (Figure 4 D, F, I). PD2 to Ach, but not carbachol is lower compared to the other segments (Figure 4 G, J) indicative of larger acetylcholinesterase activity. The mid-colon segments showed an intermediate contractile response, similar to the proximal segment but with lower spontaneous frequencies. Due to these variations, we conducted an analysis of differences between groups within each segment.

**Figure 4.**
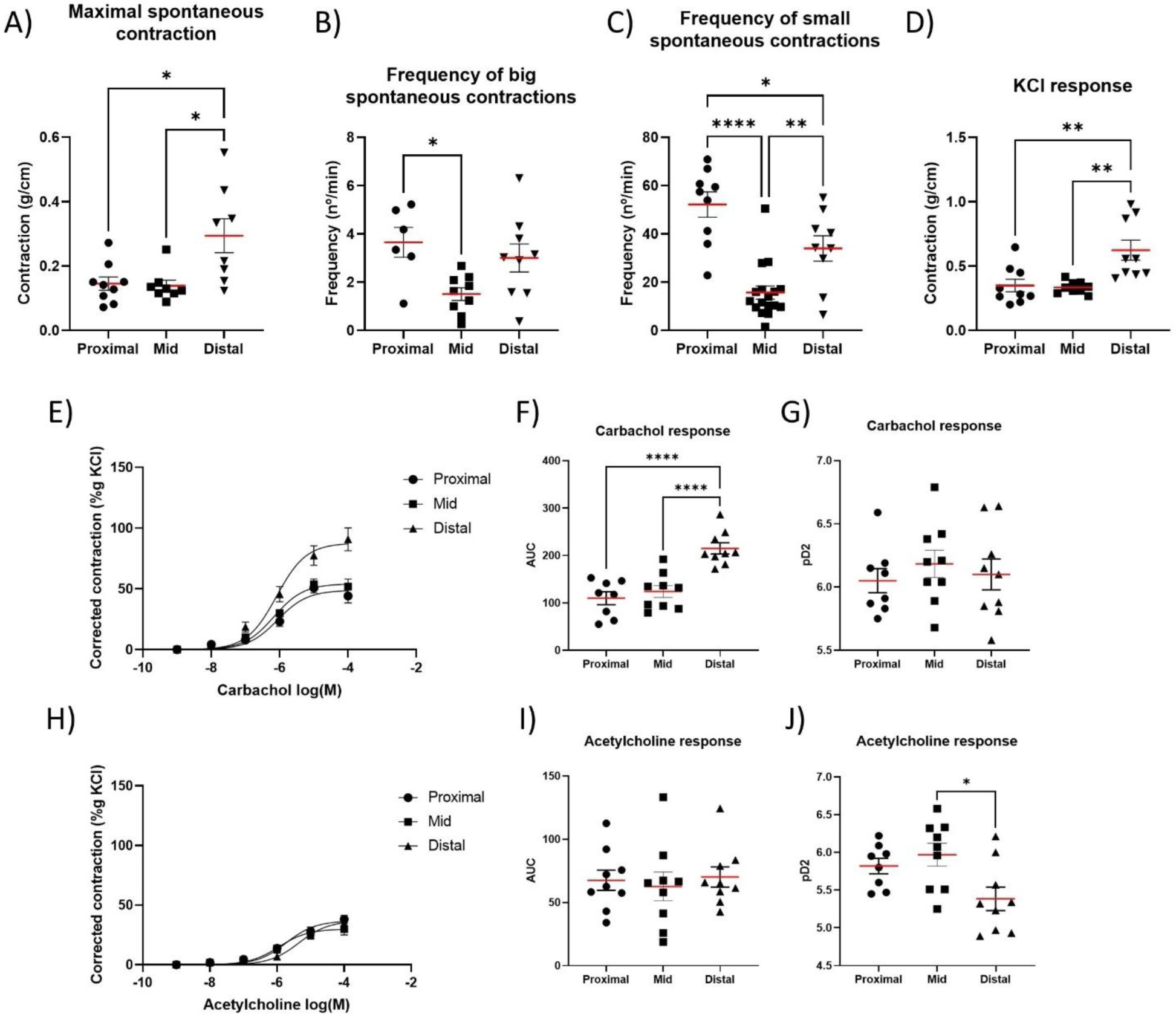
Comparison of WT mice colon segments (proximal, mid and distal) behavior in the different parameters evaluated in the organ bath system. A) Maximum of spontaneous contractions, B) frequency of big spontaneous contractions, C) frequency of small spontaneous contractions, D) maximum contractile response to KCl, E-J) contraction in response to increasing concentrations of muscarinic agonists carbachol and acetylcholine. One-way ANOVA (n = 9, * p < 0.05, ** p < 0.01, *** p < 0.001, **** p < 0.0001.

In summary, maximal spontaneous contraction and response to stimuli increase from ileum to distal colon, while the frequency of spontaneous contractions is reduced, associated to the different physiological functions during digestion.

### The absence of adaptive lymphocytes increases contractile response in the proximal colon, while colitis reduces it

In the proximal segment of the colon, closest to the ileum, we primarily observed two phenomena. Under healthy conditions, Rag1^-/-^ mice exhibited higher maximum and frequency in spontaneous contractions (Figure 5 A and B), as well as in response to KCl, compared to WT (Figure 5 D), with no differences in the AUC to carbachol or ACh between genotypes (Figure 5 F, I). However, acetylcholine PD2 was lower in Rag1^-/-^ mice compared to WT, suggesting that WT animals have higher sensitivity to ACh in their proximal colon (Figure 5 J). Since this phenomenon did not occur with carbachol (Figure 5 G), it may be due to differences in acetylcholinesterase activity, which may be increased in Rag1^-/-^ mice.

**Figure 5.**
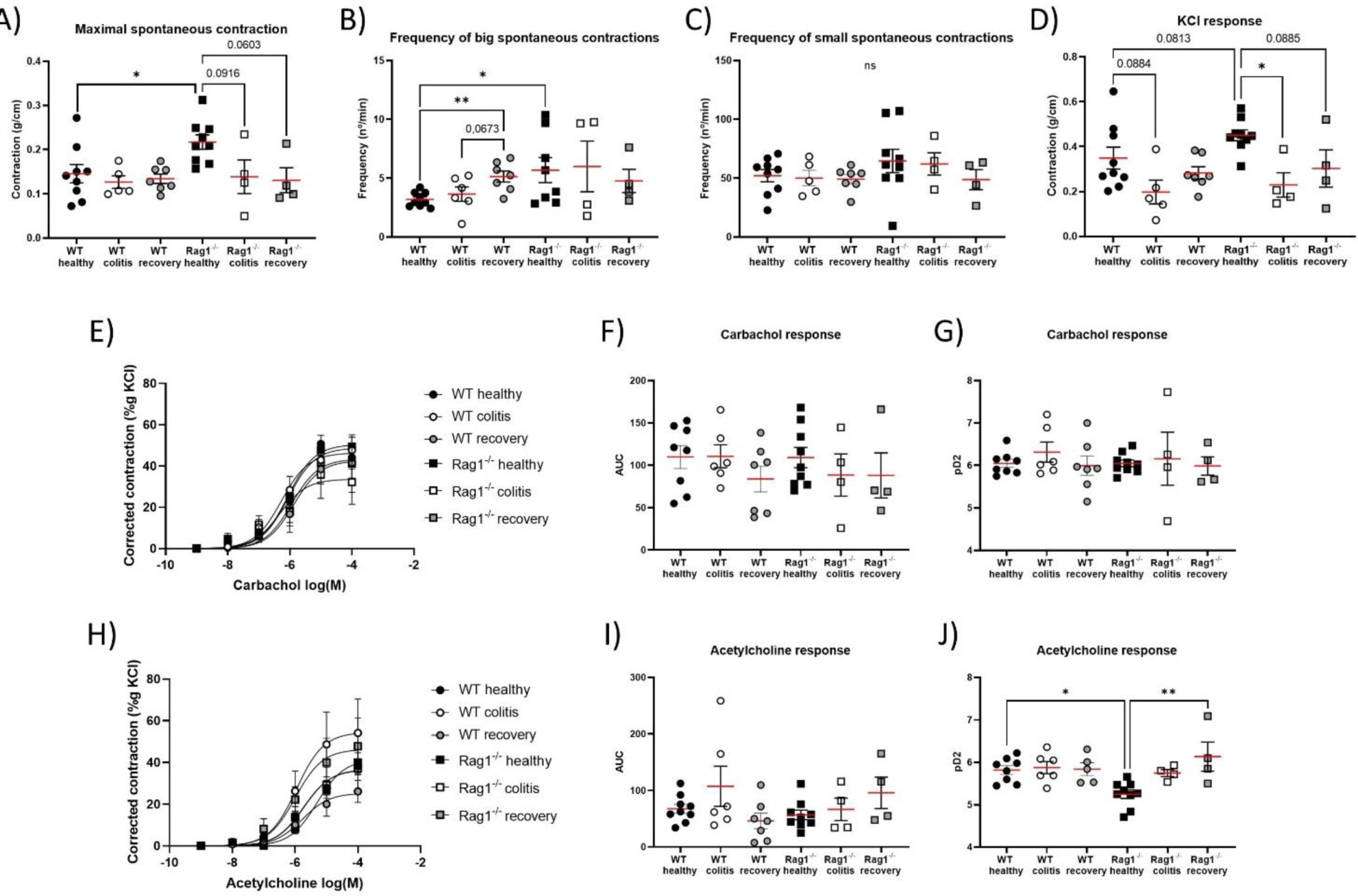
Data analysis of proximal colon segment. A) Maximum of spontaneous contractions, B) frequency of big spontaneous contractions, C) frequency of small spontaneous contractions, D) maximum contractile response to KCl, E-J) contraction in response to increasing concentrations of muscarinic agonists carbachol and acetylcholine. One-way ANOVA (n = 4 - 9) * p < 0.05, ** p < 0.01.

When assessing the effects of colitis on the spontaneous maximal contraction of the colon (Figure 5 A), colitis WT mice did not exhibit a significant difference compared to healthy WT mice. However, colitis reduced spontaneous maximal contraction in colitis Rag1^-/-^ mice compared to healthy Rag1^-/-^ mice to similar levels as in colitis WT mice. Regarding the KCl response (Figure 5 D), a similar trend was observed, with a tendency or a significant reduction in KCl response in colitis WT and colitis Rag1^-/-^ mice compared to their respective healthy groups. Carbachol and ACh responses were not affected by colitis in both WT and Rag1^-/-^ mice.

Upon recovery from colitis, no significant differences were observed in the recovery groups compared to colitis groups in either WT or Rag1^-/-^ samples in all the quantified parameters (Figure 5 A-J).

Under healthy conditions, the frequency of large contractions was higher in Rag1^-/-^ compared to WT. However, in mice with colitis, no differences were found between genotypes and when comparing colitis with healthy conditions. There were also no differences observed in the frequency of spontaneous small contractions between genotypes and between healthy and colitis conditions (Figure 5 C). Upon recovery, a significant increase in large contractions was found in WT mice, an effect that was not observed in Rag1^-/-^.

In summary, in the proximal colon, under healthy conditions the adaptive immune system seems to negatively modulate frequency and contractile response, and thus the degree of excitability. Colitis in this region does not significantly alter the frequency of contractions but does reduce contractile capacity, while recovery may increase frequency of spontaneous contractions. The adaptive immune system does not seem to play a role under colitis conditions.

### Frequency of spontaneous contractions and the contractile response of mid-colon are affected by the absence of adaptive lymphocytes during colitis but not in healthy state

We defined the mid colon as comprising the two middle segments of the four that we dissected. In healthy animals, there are no significant differences due to genotype in spontaneous contractions maximum or frequency (Figure 6 A-C), nor in the maximal response to KCl or muscarinic agonists (Figure 6 D, F, I). The only difference is a tendency of less sensibility to Ach in Rag1^-/-^ healthy mice (Figure 6 J), as observed in ileum and proximal colon.

**Figure 6.**
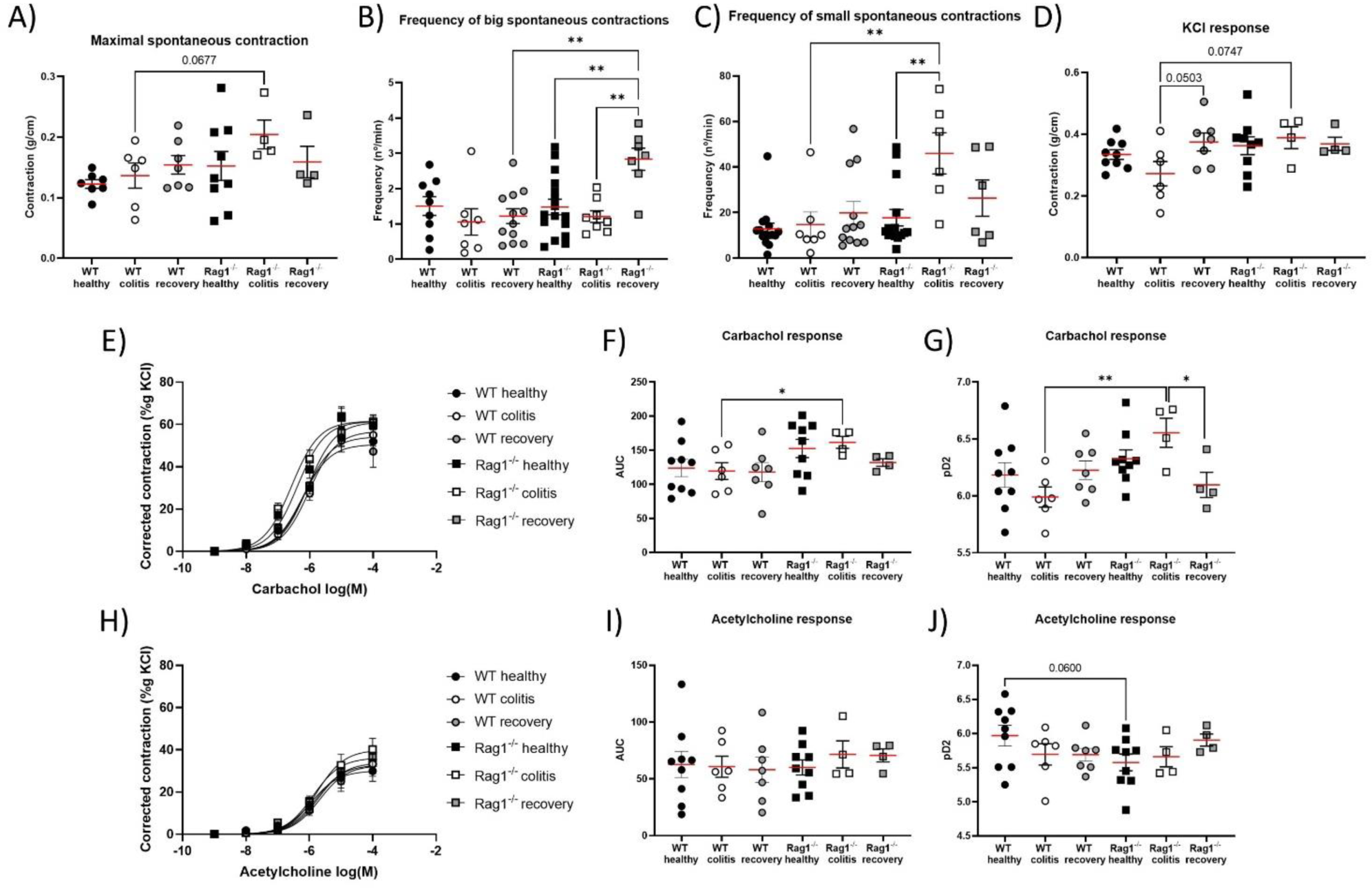
Data analysis of mid colon segment. A) Maximum of spontaneous contractions, B) frequency of big spontaneous contractions, C) frequency of small spontaneous contractions, D) maximum contractile response to KCl, E-J) contraction in response to increasing concentrations of muscarinic agonists carbachol and acetylcholine. One-way ANOVA (n = 4 - 18) * p < 0.05, ** p < 0.01.

In WT animals, no changes due to colitis were observed in any parameter, except of a tendency of less contractile response to KCl of colitis group compared to recovery. However, Rag1^-/-^ animals in colitis state, compared to WT colitis, exhibited higher contractile capacity in both spontaneous contractions and in response to KCl and carbachol (Figure 6 A, D, F), and higher sensibility to carbachol (Figure 6 G).

Distinct differences emerged in the frequency of spontaneous contractions. In WT mice, no differences were observed between healthy, colitis and recovery samples. However, in Rag1^-/-^ mice, the frequency of large contractions was higher in the recovery group (Figure 6 B), while the frequency of small contractions is higher in the colitis group (Figure 6 C).

In summary, in the mid-colon, the absence of the adaptive immune system in healthy conditions enhances the response to muscarinic agonists, while colitis has no notable impact on the frequency of contractions or contractile capacity. However, under inflammatory state of colitis, the presence of adaptive lymphocytes seems to negatively modulate the frequency and maximum of spontaneous and KCl contractions, and thus affecting the excitability degree of SMC and/or ICC, and decreasing SMC sensitivity to muscarinic agonists.

### Distal colon contraction is influenced by both colitis and the presence of adaptive lymphocytes

In the distal segment of the colon, in healthy animals, no differences are observed in maximum or frequency of spontaneous contractions between Rag1^-/-^ and WT mice (Figure 7 A-C). However, KCl contraction was significantly lower in Rag1^-/-^ without any change in the response to muscarinic agonists (Figure 7 D, F, I).

**Figure 7.**
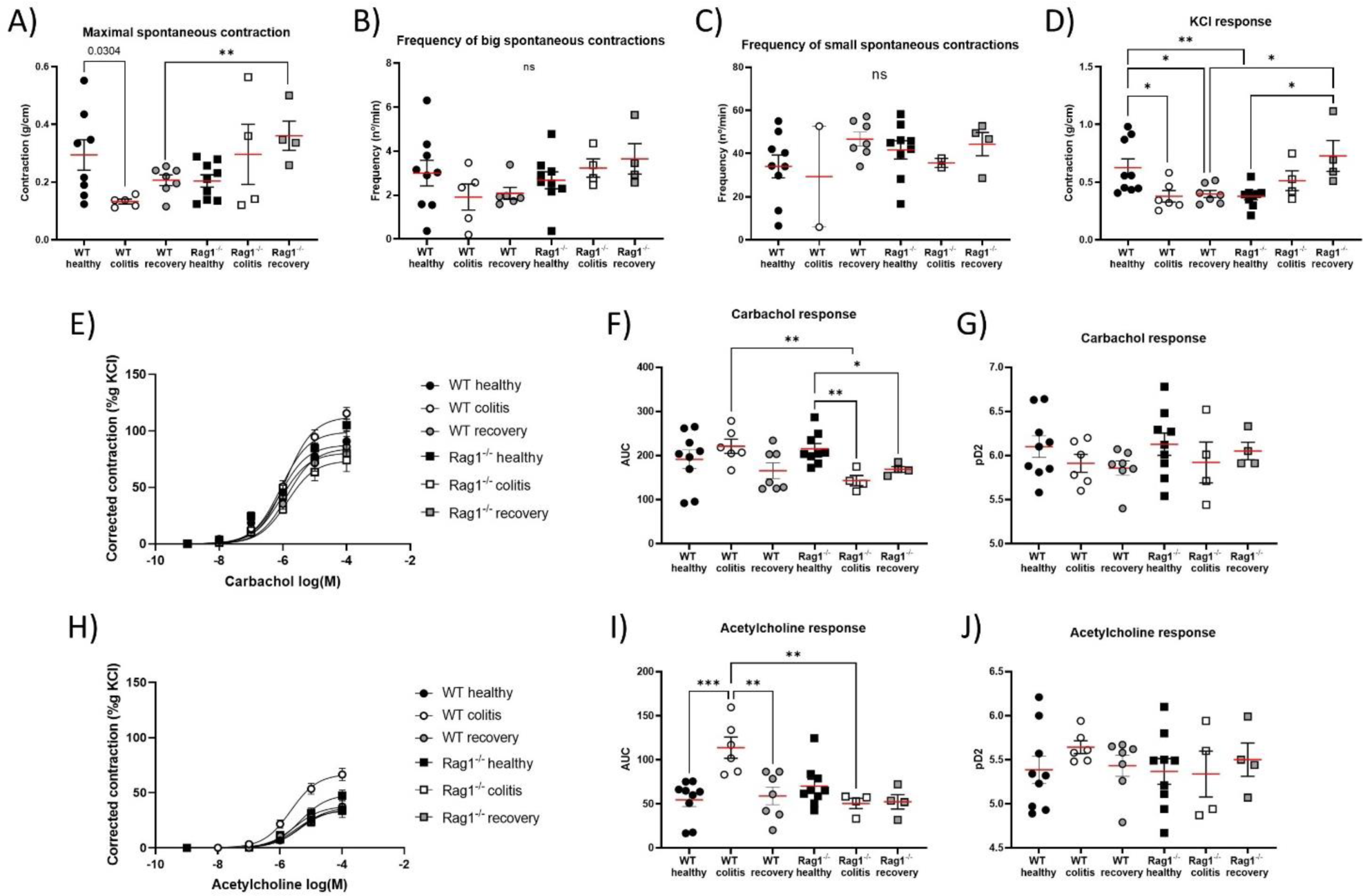
Data analysis of distal colon segment. A) Maximum of spontaneous contractions, B) frequency of big spontaneous contractions, C) frequency of small spontaneous contractions, D) maximum contractile response to KCl, E-J) contraction in response to increasing concentrations of muscarinic agonists carbachol and acetylcholine. One-way ANOVA (n = 4 - 9) * p < 0.05, ** p < 0.01.

In WT animals, colitis induces a decrease in contractile capacity, as seen in both spontaneous maximal contractions (Figure 7 A) and responses to KCl (Figure 7 D) compared to healthy WT. Interestingly, the colitis WT group exhibits a heightened contractile response to Ach (Figure 7 I), while the response to carbachol remains unchanged by colitis (Figure 7 E-G). Conversely, Rag1^-/-^ mice show the opposite behavior, with a trend towards increased contractile capacity in the colitis group and a significant increase in the recovery group in spontaneous contractions and in response to KCl (Figure 7 A, D), and a decreased response to carbachol that is maintained during recovery (Figure 7 F).

Regarding the frequency of spontaneous contractions, no changes were observed in either large or small contractions among groups (Figure 7 B and C).

In summary, in the distal colon under healthy conditions, the adaptive immune system enhances KCl response without modification of muscarinic contractions, suggesting a positive modulation on SMC excitability. Under colitis conditions, the presence of adaptive lymphocytes enhances Ach and carbachol responses, suggesting, that under an inflammatory environment they increase the response of muscarinic receptors, which may contribute to enhanced motility.

## Discussion

Our findings suggest that the adaptive immune system regulates intestinal motility in both healthy conditions and during colitis. In this study, we aimed to investigate the role of adaptive immune system on contractility of different segments of the small and large intestine and the modulation T and B lymphocytes exert in the context of IBD. Although IBD is considered as an innate-mediated inflammation, T cells play an important role ^7-9^, so in this study, we decided to examine the impact of adaptive immune cells and inflammation on the characteristics of ileal and colonic motility using Rag1^-/-^ mice lacking adaptive immune cells (T and B cells)^16^.

Intestinal motility is regulated mainly by slow waves produced by ICCs, muscle distension and neurotransmitters such as Ach coming from myenteric plexus ^20^. Gastrointestinal motor functions depend on its two layers of smooth muscle: circular and longitudinal, which function in a coordinated manner, and which are excited by two basic types of electrical waves: 1) slow waves, which condition the rhythm of contractions and are possibly by the ICCs, and 2) spikes, which are true action potentials that occur when the slow waves exceed the -40mV threshold. Muscle distension and neurotransmitters such as acetylcholine coming from the myenteric plexus depolarize the SMC and make them more excitable ^20^. We, therefore, evaluated different aspects of motility, including frequency of spontaneous contractions, which depend on ICC activity, contractility to KCl which is regulated by the depolarizing state of the SMC, and the responses to ACh and carbachol. ACh is recognized as a primary neurotransmitter involved in the intestinal smooth muscle contraction, acting through M2 and M3 muscarinic receptors, and its activity is also regulated by the acetylcholinesterase, an enzyme responsible for ACh degradation ^13,20^. The study of both agonists allowed us to examine possible modifications on muscarinic receptor sensitivity and acetylcholinesterase activity.

It is worth noting that while previous studies have mainly focused on circular smooth muscle contractions, our system allows for the investigation of longitudinal smooth muscle contractions due to the preparation’s assembly. We analysed different regions, including the ileum and three regions of the colon, since they have different role. The main function of the ileum is to absorb nutrients derived from the digestion of food that have not been absorbed in the anterior regions of the small intestine, while the colon has two main functions: absorbing water, electrolytes and metabolites derived from the microbiota (mainly proximal colon), and storing fecal matter until its expulsion (distal colon). The colonic motor cycle consists of two stages, quiescent and contractile stages, whereas a typical motor cycle in the small intestine involves a series of stages starting from a quiescent state, followed by progressively increasing amplitude and frequency of contractions, leading to a state of maximal contractile activity ^21^.

In healthy mice under physiological conditions, adaptive immune cells help to maintain a low contractile response and frequency in proximal colon, where water, electrolytes and nutrient derived from the microbiota are primarily absorbed ^20^. This allows for sufficient time for the absorption process ^22,23^. However, in an inflammatory environment, such as acute colitis, significant changes in contractile response are observed. In colitis, adaptive immune cells promote a high frequency of spontaneous contractions in ileum and proximal colon, an effect that is reverted in mid and distal colon. Additionally, the strength of spontaneous contractions in the ileum is enhanced. However, in the colon, the inflammation coupled with adaptive immune activity reduces the contractile capacity of colon walls, particularly in the mid and distal colon. Thus, it could be that the adaptive immune system is accelerating the transit in the nutrient absorption zone but retaining its advance prior to defecation ^20^.

Our findings suggest that inflammation and the presence of adaptive immune cells can alter the response to muscarinic agonists, whether through muscarinic receptors or acetylcholinesterase activity. In ileum, proximal and mid colon, our data suggested that the acquired immune system in healthy conditions participates in the contractile effect of acetylcholine, as its lack reduced PD2 values. These data indicate that adaptive lymphocytes either increase muscarinic receptor sensitivity or reduce acetylcholinesterase activity. Since the effect was not observed with carbachol, we suggest that, under healthy conditions, the adaptive immune system may reduce acetylcholinesterase activity and, thus, prolong the action of acetylcholine, rather than affecting the receptors themselves. On the other hand, in ileum and mid colon, in the presence of inflammation associated with DSS treatment, WT mice presented reduced PD2 compared to healthy mice to Ach, but higher PD2 values to carbachol in colitis Rag1^-/-^ mice. These data suggest that in a situation of noxious stimuli, the adaptive immune system, or associated inflammation, reduces the sensibility of the muscarinic receptors. This effect seems to be maintained during recovery in ileum with fewer muscarinic receptors (less contractile response to both carbachol and Ach) but increased sensitivity (a trend to higher PD2 values to carbachol). Lastly, in the distal colon, adaptive immunity in colitis reduced contractile capacity of spontaneous contractions and KCl depolarization but increased the contractile response to muscarinic agonists without affecting the PD2, which again suggests a reduction in the activity of acetylcholinesterase. Several studies indicate that high levels of acetylcholine act as anti-inflammatory substance in colitis, pointing to acetylcholinesterase inhibition as a possible treatment ^24,25^. Here, we have seen that adaptive immune system in healthy conditions is probably reducing acetylcholinesterase activity from ileum to mid colon, protecting from inflammation. In the context of intestinal inflammation, studies have shown changes in muscarinic receptor activity. For example, in a dog ileitis model induced by acetic acid, alterations in muscarinic receptor function were observed. Specifically, while the contraction of circular smooth muscle in response to acetylcholine decreased, the contraction of longitudinal smooth muscle remained unchanged ^26^. In a guinea-pig ileitis model induced by trinitrobenzene sulphonic acid (TNBS), the contractile force of the circular smooth muscle decreases in response to acetylcholine, while the contractile force of the longitudinal smooth muscle increases ^27^. On the other hand, some reports suggest that TNBS-induced inflammation reduces muscarinic receptor-mediated contraction of longitudinal smooth muscle ^13,28^. Our experimental approach using an organ bath allows for a detailed examination of the contractile response of the longitudinal smooth muscle. Our results in ileum suggest that colitis reduces sensibility of muscarinic receptors mediated by adaptive immune cells or inflammation-derived substances, but we also observed a higher response to Ach in distal colon, probably due to low acetylcholinesterase activity during colitis.

Changes in the strength of spontaneous contractions and in the response to KCl can be explained by possible changes in the depolarization of the cells, as wells as the frequency of spontaneous contractions, which also can be affected by the functioning of ICC. For example, in the proximal colon of healthy mice, the absence of adaptive immune system increases spontaneous contractions and KCl contractions, with no effect on maximal muscarinic agonists-induced contractions, suggesting possible changes in the depolarization level of SMCs. The adaptive immune system may have a depressant effect on colonic contractions under healthy conditions, modifying the depolarization level of the SMC, possibly though modulation of membrane channels. The higher frequency of spontaneous contractions in Rag1^-/-^ mice may be related to modifications in ICCs by the adaptive immune system (the presence of adaptive lymphocytes could reduce firing rate of depolarization) or could also act at the channel level. In the presence of a noxious stimuli, such as DSS, a larger effect was observed on Rag1^-/-^ mice, lowering the strength of spontaneous and KCl contractions and frequency (i.e. in the presence of noxious stimuli, the adaptive immune system may contribute to increase colonic contractions), both in ileum and proximal colon, while the effect is inverted in the medium and distal colon. Previous research has shown that inflammation has a significant impact on gastrointestinal motility patterns ^13,29^, resulting in decreased excitability of SMC^29^. This effect is particularly evident in the distal colon, as observed in our study. However, the extent of this effect appears to vary depending on the type of SMC involved, circular versus longitudinal, and the specific experimental model employed ^30^.

Emerging evidence suggests that inflammation not only alters the expression of ion channels but also affects the enteric nervous system and the function of SMC ^13,29^.

Another possibility explaining the reduction in contractile responses under inflammatory conditions observed in our study during acute colitis could be an upregulation of K^+^ channels, or a decrease in Ca^2+^ channels, as suggested by several studies ^29^. Various inflammatory models exhibit a reduction in the amplitude of Ca^2+^ currents in SMC as observed by patch-clamp recordings in murine DSS-induced colitis and ethanol/acetic acid-induced inflammation in canine colon, in association with a decrease in expression in pore-forming α Ca2+ channel subunit ^31,32^. In TNBS-induced inflammation of the rat colon, Ca^2+^ currents also decrease together with attenuated responses to Ca2+ channel agonists, which may be regulated by auxiliary subunits that influence channel activation kinetics ^33^, with nuclear factor (NF)-κB being the transcriptional factor involved in the expression these genes. It is worth noting that NF-κB upregulation has been reported in patients with inflammatory bowel disease ^34,35^, with enhanced DNA binding in smooth muscle from inflamed colon ^36^. However, inflammation has a selective impact on membrane ion channels. In DSS mice, Ca2+ currents were downregulated, while the amplitude and kinetics of the transient outward K+ current remained unaltered ^31^. However, adenosine triphosphate (ATP)-sensitive K+ channel (KATP) exhibited increased bursting behavior and heightened sensitivity to the channel opener ^29,31^. It is still unclear whether these alterations are due to cells of the adaptive immune system, and further studies in more comprehensive models, rather than isolated muscle cells, are required to elucidate the underlying cause.

Differences in cytokine profiles can indeed account for variations in contractile responses. For example, in a model of gut inflammation induced by nematode infection, such as *Trichinella spiralis* infection, smooth muscle contractility is increased. ^37,38^. Conversely, smooth muscle contractility is reduced in cases of intestinal inflammation induced by TNBS, surgical manipulation, experimental obstruction, hemorrhagic shock, or peritonitis. ^13,28,39-44^. According to reports from the Collins group ^45-47^, hypercontractility induced by nematodes is mediated by an increase in prostaglandin E2 (PGE2) levels following the induction of Th2 cytokines such as interleukin (IL)-4 and IL-13. Conversely, TNBS-induced gut inflammation is mainly mediated by Th1 cytokines, such as IL-1β, tumor necrosis factor-α (TNF-α), and IL-12 ^13,41,42,48^. Our model examines three situations characterized by different cytokine profiles. In healthy mice, a presumed homeostatic situation exists, with a low number of immune cells producing mainly regulatory molecules such as IL-10, IL-35, and TGF-β. These cells include Tregs, some Th17, and other innate immune cells such as innate lymphoid cells or myeloid cells (dendritic cells, macrophages) ^7-9^. During the acute phase of DSS-induced colitis, similar to the TNBS model, intestinal inflammation is mediated by type 1 cytokines produced by Th1 and other pro-inflammatory cells ^7-9^. In the recovery phase of colitis, inflammation is reduced, and regulatory cells attempt to restore normality, although immune cell infiltration persists. In WT animals, the transition to a Th1 pro-inflammatory phenotype during colitis increases the frequency of spontaneous contractions and the contractile capacity in the ileum and decreases them in the colon. And in the response to muscarinic agonists, adaptive immune cells in homeostasis, mainly Treg, reduce acetylcholinesterase activity increasing the sensibility to Ach, while Th1 and the proinflammatory ambient during acute inflammation reduces the sensibility of muscarinic receptors in ileum but increases Ach response in distal colon by reducing acetylcholinesterase activity. These changes largely return to normal during recovery, except for a decreased muscarinic response in ileum due to low number of receptors, and increase in spontaneous contraction frequency in the proximal colon and a persistent decrease in contractile response in the distal colon. Thus, B and T cells, particularly Treg cells, may contribute to the maintenance of a homeostatic motility phenotype and a low acetylcholinesterase activity that protects from inflammation ^24,25^, while during colitis, Th1 cells primarily increase the frequency and contractile response in the intestinal highways, particularly in the ileum, while decreasing it along the colonic segments, and reduces muscarinic response in ileum but increases it in distal colon.

Our findings demonstrate that both colitis and adaptive immune system significantly impact intestinal motility in both the ileum and colon. Specifically, we observed that in healthy animals, the adaptive immune system, likely through Tregs, regulates acetylcholine contractions, probably by maintaining a low acetylcholinesterase activity, which may have a protective role. In colitis, the inflammation reduced contractile capacity and frequency of contractions, which returned to normal levels upon recovery. Additionally, the presence of adaptive immune cells has a negative impact on the length of ileum after DSS treatment, suggesting a role in the pathology. Based on the data from other models of intestinal inflammation we speculate that this may be related to changes in Ca^2+^ channels and/or nerve damage, and somehow adaptive immune cells present in this inflammatory ambient, Th1 mainly, may contribute. Unravelling these mechanisms and how adaptive immune cells and their associated cytokine milieu influence motility requires further investigation.

## Conclusions

In conclusion, this study reveals alterations in intestinal motility during IBD, emphasizing the role of adaptive immunity. The organ bath system proves valuable in segment-specific analyses, shedding light on distinct responses in the ileum and various colon segments. Understanding these motility changes contributes to the broader understanding of IBD pathology and highlights the need for comprehensive investigations beyond inflammation. This exploration provides a foundation for future studies aiming to address the intricacies of gastrointestinal motility in health and disease.

## Funding

This research was funded by Ministerio de Ciencia, Innovación y Universidades from Spain (grant number RTI2018-097504-B-I00), Instituto de Salud Carlos III (ISCIII) (grant number PI20/00306) with co-funding from the European Regional Development Fund (ERDF) “A way to build Europe”. RGB was supported by the UAM and the MCNU FPU program (FPU19/01774) and AS by Universidad Francisco de Vitoria.

## Author Contributions

R.G.B, Data curation, Formal Analysis, Investigation, Methodology. Software, Validation, Visualization, Writing – original draft, Writing-review & editing. P.R.R.: Data curation, Formal Analysis, Investigation, Methodology. Writing – review & editing. M.O.Z, S.R., A.S.: Formal Analysis, Investigation, Methodology, Visualization, Writing-review & editing. S.M.A.: Conceptualization, Data curation, Formal Analysis, Funding acquisition, Investigation, Methodology. Project administration, Resources, Software, Supervision, Validation, Visualization, Writing-original draft, Writing-review & editing. J.M.G.G. Conceptualization, Funding acquisition, Investigation, Project administration, Resources, Software, Supervision, Validation, Visualization, Writing-original draft, Writing-review & editing.

## Competing interest

There are no potential conflicts of interest.

## Abbreviations

Ach: Acetylcholine
AUC: Area under the curve
CD: Crohn’s Disease
DCs: Dendritic cells
DSS: Dextran sulfate sodium
ENS: Enteric nervous system
IBD: Inflammatory bowel disease
IBS: Irritable bowel syndrome
ICCs: Interstitial cells of Cajal
IL: Interleukin
KATP: Adenosine triphosphate (ATP)-sensitive K+ channel
NF-Κb: Nuclear factor κB
PD2: -logEC50
PGE2: Prostaglandin E2
SMCs: Smooth muscle cells
SPF: Specific pathogen-free
TNF-α: tumor necrosis factor-α
Th: T helper cell
Tregs: Regulatory T cells
UC: Ulcerative Colitis
WT: Wild-type

## Notes

### Competing Interest Statement

The authors have declared no competing interest.

## References

1. Li, N., Zuo, B., Huang, S., Zeng, B., Han, D., Li, T., Liu, T., Wu, Z., Wei, H., Zhao, J., and Wang, J. (2020). Spatial heterogeneity of bacterial colonization across different gut segments following inter-species microbiota transplantation. Microbiome 8, 161. 10.1186/s40168-020-00917-7.

2. Mowat, A.M., and Agace, W.W. (2014). Regional specialization within the intestinal immune system. Nat Rev Immunol 14, 667–685. 10.1038/nri3738.

3. Fenton, T.M., Jorgensen, P.B., Niss, K., Rubin, S.J.S., Morbe, U.M., Riis, L.B., Da Silva, C., Plumb, A., Vandamme, J., Jakobsen, H.L., et al. (2020). Immune Profiling of Human Gut-Associated Lymphoid Tissue Identifies a Role for Isolated Lymphoid Follicles in Priming of Region-Specific Immunity. Immunity 52, 557–570 e556. 10.1016/j.immuni.2020.02.001.

4. Villablanca, E.J., Wang, S., de Calisto, J., Gomes, D.C., Kane, M.A., Napoli, J.L., Blaner, W.S., Kagechika, H., Blomhoff, R., Rosemblatt, M., et al. (2011). MyD88 and retinoic acid signaling pathways interact to modulate gastrointestinal activities of dendritic cells. Gastroenterology 141, 176–185. 10.1053/j.gastro.2011.04.010.

5. Parigi, S.M., Larsson, L., Das, S., Ramirez Flores, R.O., Frede, A., Tripathi, K.P., Diaz, O.E., Selin, K., Morales, R.A., Luo, X., et al. (2022). The spatial transcriptomic landscape of the healing mouse intestine following damage. Nat Commun 13, 828. 10.1038/s41467-022-28497-0.

6. Brazil, J.C., Quiros, M., Nusrat, A., and Parkos, C.A. (2019). Innate immune cell-epithelial crosstalk during wound repair. The Journal of clinical investigation 129, 2983–2993. 10.1172/JCI124618.

7. Gomez-Bris, R., Saez, A., Herrero-Fernandez, B., Rius, C., Sanchez-Martinez, H., and Gonzalez-Granado, J.M. (2023). CD4 T-Cell Subsets and the Pathophysiology of Inflammatory Bowel Disease. Int J Mol Sci 24. 10.3390/ijms24032696.

8. Saez, A., Gomez-Bris, R., Herrero-Fernandez, B., Mingorance, C., Rius, C., and Gonzalez-Granado, J.M. (2021). Innate Lymphoid Cells in Intestinal Homeostasis and Inflammatory Bowel Disease. Int J Mol Sci 22. 10.3390/ijms22147618.

9. Saez, A., Herrero-Fernandez, B., Gomez-Bris, R., Sanchez-Martinez, H., and Gonzalez-Granado, J.M. (2023). Pathophysiology of Inflammatory Bowel Disease: Innate Immune System. Int J Mol Sci 24. 10.3390/ijms24021526.

10. Liu, D., Saikam, V., Skrada, K.A., Merlin, D., and Iyer, S.S. (2022). Inflammatory bowel disease biomarkers. Med Res Rev 42, 1856–1887. 10.1002/med.21893.

11. Huizinga, J.D., and Lammers, W.J. (2009). Gut peristalsis is governed by a multitude of cooperating mechanisms. Am J Physiol Gastrointest Liver Physiol 296, G1–8. 10.1152/ajpgi.90380.2008.

12. Seerden, T.C., Lammers, W.J., De Winter, B.Y., De Man, J.G., and Pelckmans, P.A. (2005). Spatiotemporal electrical and motility mapping of distension-induced propagating oscillations in the murine small intestine. Am J Physiol Gastrointest Liver Physiol 289, G1043–1051. 10.1152/ajpgi.00205.2005.

13. Ohama, T., Hori, M., and Ozaki, H. (2007). Mechanism of abnormal intestinal motility in inflammatory bowel disease: how smooth muscle contraction is reduced? J Smooth Muscle Res 43, 43–54. 10.1540/jsmr.43.43.

14. Bassotti, G., Antonelli, E., Villanacci, V., Salemme, M., Coppola, M., and Annese, V. (2014). Gastrointestinal motility disorders in inflammatory bowel diseases. World J Gastroenterol 20, 37–44. 10.3748/wjg.v20.i1.37.

15. Khan, W.I., Richard, M., Akiho, H., Blennerhasset, P.A., Humphreys, N.E., Grencis, R.K., Van Snick, J., and Collins, S.M. (2003). Modulation of intestinal muscle contraction by interleukin-9 (IL-9) or IL-9 neutralization: correlation with worm expulsion in murine nematode infections. Infection and immunity 71, 2430–2438. 10.1128/IAI.71.5.2430-2438.2003.

16. Mombaerts, P., Iacomini, J., Johnson, R.S., Herrup, K., Tonegawa, S., and Papaioannou, V.E. (1992). RAG-1-deficient mice have no mature B and T lymphocytes. Cell 68, 869–877. 10.1016/0092-8674(92)90030-g.

17. Saiz, M.L., Cibrian, D., Ramirez-Huesca, M., Torralba, D., Moreno-Gonzalo, O., and Sanchez-Madrid, F. (2017). Tetraspanin CD9 Limits Mucosal Healing in Experimental Colitis. Front Immunol 8, 1854. 10.3389/fimmu.2017.01854.

18. Jespersen, B., Tykocki, N.R., Watts, S.W., and Cobbett, P.J. (2015). Measurement of smooth muscle function in the isolated tissue bath-applications to pharmacology research. J Vis Exp, 52324. 10.3791/52324.

19. Instruments, A. Isolated Tissue Baths. https://www.adinstruments.com/research/in-vitro/pharmacology-isolated-tissue-and-organs/isolated-tissue-baths.

20. E. Hall, J. (2011). Guyton and Hall Textbook of Medical Physiology (Elsevier).

21. Sarna, S.K. (1985). Cyclic motor activity; migrating motor complex: 1985. Gastroenterology 89, 894–913. 10.1016/0016-5085(85)90589-x.

22. Treichel, A.J., Finholm, I., Knutson, K.R., Alcaino, C., Whiteman, S.T., Brown, M.R., Matveyenko, A., Wegner, A., Kacmaz, H., Mercado-Perez, A., et al. (2022). Specialized Mechanosensory Epithelial Cells in Mouse Gut Intrinsic Tactile Sensitivity. Gastroenterology 162, 535–547 e513. 10.1053/j.gastro.2021.10.026.

23. Kiela, P.R., and Ghishan, F.K. (2016). Physiology of Intestinal Absorption and Secretion. Best Pract Res Clin Gastroenterol 30, 145–159. 10.1016/j.bpg.2016.02.007.

24. Singh, S.P., Chand, H.S., Banerjee, S., Agarwal, H., Raizada, V., Roy, S., and Sopori, M. (2020). Acetylcholinesterase Inhibitor Pyridostigmine Bromide Attenuates Gut Pathology and Bacterial Dysbiosis in a Murine Model of Ulcerative Colitis. Dig Dis Sci 65, 141–149. 10.1007/s10620-019-05838-6.

25. Zheng, W., Song, H., Luo, Z., Wu, H., Chen, L., Wang, Y., Cui, H., Zhang, Y., Wang, B., Li, W., et al. (2021). Acetylcholine ameliorates colitis by promoting IL-10 secretion of monocytic myeloid-derived suppressor cells through the nAChR/ERK pathway. Proc Natl Acad Sci U S A 118. 10.1073/pnas.2017762118.

26. Shi, X.Z., and Sarna, S.K. (1999). Differential inflammatory modulation of canine ileal longitudinal and circular muscle cells. Am J Physiol 277, G341–350. 10.1152/ajpgi.1999.277.2.G341.

27. Martinolle, J.P., Garcia-Villar, R., Fioramonti, J., and Bueno, L. (1997). Altered contractility of circular and longitudinal muscle in TNBS-inflamed guinea pig ileum. Am J Physiol 272, G1258–1267. 10.1152/ajpgi.1997.272.5.G1258.

28. Moreels, T.G., De Man, J.G., Dick, J.M., Nieuwendijk, R.J., De Winter, B.Y., Lefebvre, R.A., Herman, A.G., and Pelckmans, P.A. (2001). Effect of TNBS-induced morphological changes on pharmacological contractility of the rat ileum. Eur J Pharmacol 423, 211–222. 10.1016/s0014-2999(01)01088-3.

29. Malykhina, A.P., and Akbarali, H.I. (2004). Inflammation-induced “channelopathies” in the gastrointestinal smooth muscle. Cell Biochem Biophys 41, 319–330. 10.1385/CBB:41:2:319.

30. Imanishi, T., Hara, H., Suzuki, S., Suzuki, N., Akira, S., and Saito, T. (2007). Cutting edge: TLR2 directly triggers Th1 effector functions. J Immunol 178, 6715–6719. 10.4049/jimmunol.178.11.6715.

31. Akbarali, H.I., Pothoulakis, C., and Castagliuolo, I. (2000). Altered ion channel activity in murine colonic smooth muscle myocytes in an experimental colitis model. Biochem Biophys Res Commun 275, 637–642. 10.1006/bbrc.2000.3346.

32. Liu, X., Rusch, N.J., Striessnig, J., and Sarna, S.K. (2001). Down-regulation of L-type calcium channels in inflamed circular smooth muscle cells of the canine colon. Gastroenterology 120, 480–489. 10.1053/gast.2001.21167.

33. Kinoshita, K., Sato, K., Hori, M., Ozaki, H., and Karaki, H. (2003). Decrease in activity of smooth muscle L-type Ca2+ channels and its reversal by NF-kappaB inhibitors in Crohn’s colitis model. Am J Physiol Gastrointest Liver Physiol 285, G483–493. 10.1152/ajpgi.00038.2003.

34. Neurath, M.F., Fuss, I., Schurmann, G., Pettersson, S., Arnold, K., Muller-Lobeck, H., Strober, W., Herfarth, C., and Buschenfelde, K.H. (1998). Cytokine gene transcription by NF-kappa B family members in patients with inflammatory bowel disease. Ann N Y Acad Sci 859, 149–159. 10.1111/j.1749-6632.1998.tb11119.x.

35. Ghosh, S., May, M.J., and Kopp, E.B. (1998). NF-kappa B and Rel proteins: evolutionarily conserved mediators of immune responses. Annu Rev Immunol 16, 225–260. 10.1146/annurev.immunol.16.1.225.

36. Shi, X.Z., Lindholm, P.F., and Sarna, S.K. (2003). NF-kappa B activation by oxidative stress and inflammation suppresses contractility in colonic circular smooth muscle cells. Gastroenterology 124, 1369–1380. 10.1016/s0016-5085(03)00263-4.

37. Vermillion, D.L., and Collins, S.M. (1988). Increased responsiveness of jejunal longitudinal muscle in Trichinella-infected rats. Am J Physiol 254, G124–129. 10.1152/ajpgi.1988.254.1.G124.

38. Blennerhassett, M.G., Vignjevic, P., Vermillion, D.L., and Collins, S.M. (1992). Inflammation causes hyperplasia and hypertrophy in smooth muscle of rat small intestine. Am J Physiol 262, G1041–1046. 10.1152/ajpgi.1992.262.6.G1041.

39. Hierholzer, C., Kalff, J.C., Billiar, T.R., Bauer, A.J., Tweardy, D.J., and Harbrecht, B.G. (2004). Induced nitric oxide promotes intestinal inflammation following hemorrhagic shock. Am J Physiol Gastrointest Liver Physiol 286, G225–233. 10.1152/ajpgi.00447.2002.

40. Kalff, J.C., Schraut, W.H., Simmons, R.L., and Bauer, A.J. (1998). Surgical manipulation of the gut elicits an intestinal muscularis inflammatory response resulting in postsurgical ileus. Ann Surg 228, 652–663. 10.1097/00000658-199811000-00004.

41. Kinoshita, K., Hori, M., Fujisawa, M., Sato, K., Ohama, T., Momotani, E., and Ozaki, H. (2006). Role of TNF-alpha in muscularis inflammation and motility disorder in a TNBS-induced colitis model: clues from TNF-alpha-deficient mice. Neurogastroenterol Motil 18, 578–588. 10.1111/j.1365-2982.2006.00784.x.

42. Kiyosue, M., Fujisawa, M., Kinoshita, K., Hori, M., and Ozaki, H. (2006). Different susceptibilities of spontaneous rhythmicity and myogenic contractility to intestinal muscularis inflammation in the hapten-induced colitis. Neurogastroenterol Motil 18, 1019–1030. 10.1111/j.1365-2982.2006.00841.x.

43. Koyluoglu, G., Kaya, T., Bagcivan, I., and Yildiz, T. (2002). Effect of L-NAME on decreased ileal muscle contractility induced by peritonitis in rats. J Pediatr Surg 37, 901–905. 10.1053/jpsu.2002.32907.

44. Won, K.J., Suzuki, T., Hori, M., and Ozaki, H. (2006). Motility disorder in experimentally obstructed intestine: relationship between muscularis inflammation and disruption of the ICC network. Neurogastroenterol Motil 18, 53–61. 10.1111/j.1365-2982.2005.00718.x.

45. Akiho, H., Blennerhassett, P., Deng, Y., and Collins, S.M. (2002). Role of IL-4, IL-13, and STAT6 in inflammation-induced hypercontractility of murine smooth muscle cells. Am J Physiol Gastrointest Liver Physiol 282, G226-232. 10.1152/ajpgi.2002.282.2.G226.

46. Akiho, H., Deng, Y., Blennerhassett, P., Kanbayashi, H., and Collins, S.M. (2005). Mechanisms underlying the maintenance of muscle hypercontractility in a model of postinfective gut dysfunction. Gastroenterology 129, 131–141. 10.1053/j.gastro.2005.03.049.

47. Akiho, H., Lovato, P., Deng, Y., Ceponis, P.J., Blennerhassett, P., and Collins, S.M. (2005). Interleukin-4- and -13-induced hypercontractility of human intestinal muscle cells-implication for motility changes in Crohn’s disease. Am J Physiol Gastrointest Liver Physiol 288, G609–615. 10.1152/ajpgi.00273.2004.

48. Neurath, M.F., Fuss, I., Kelsall, B.L., Stuber, E., and Strober, W. (1995). Antibodies to interleukin 12 abrogate established experimental colitis in mice. J Exp Med 182, 1281–1290. 10.1084/jem.182.5.1281.

